# The Genome of the Wasp *Anastatus disparis* Reveals Energy Metabolism Adaptations for Extreme Aggression

**DOI:** 10.1101/2022.10.10.511560

**Authors:** Pengcheng Liu, Ziyin Wang, Yumei Tao, Siyu Yin, Jianrong Wei, Jianjun Wang, Haoyuan Hu

**Author notes:** Corresponding authors., (Liu P), (Hu H).

## Abstract

Extremely aggressive behavior is rare in most species, as contestants can be severely injured or killed. Such high level of aggression can evolve when critical resources are limited, as the benefits of winning outweigh the potential costs of conflict. Currently, studies of extreme aggression are mainly from the perspectives of behavioral ecology and evolutionary biology, displaying distinct results from common nonextreme aggression. Here, we provide a high-quality genome of the generalist endoparasitoid *Anastatus disparis*, the males of which exhibit extreme mate-competition aggression, to shed light on possible genomic adaptations for extreme aggressive behavior. We combined Nanopore PromethION sequencing with Hi-C technology to assemble a high-quality chromosome-level genome of *A. disparis*. The size of the genome of this species (939.58 Mb) is larger than that of most hymenopterans (ranging from 180 Mb to 340 Mb) due to the expansion of repeated sequences (612.90 Mb, 65.23% of the whole genome). With the aid of RNA sequencing, 19,246 protein-coding genes were identified, and a great expansion of genes involved in detoxification was detected, which could represent an adaptation of this species to exploit a diverse range of known hosts. The integrated multiomics analysis highlighted genes involved in energy metabolism (especially from lipids) and antibacterial activity, both of which are possibly major aspects of adaptation for extreme aggression in *A. disparis*. Our study provides insight into molecular and evolutionary studies of extreme aggression in *A. disparis* and provides a valuable genomic resource for further research into the molecular basis of trait evolution in Hymenoptera.

## Introduction

Aggressive behavior is important for animal survival and reproduction. The acquisition of limited resources (*i.e.*, food and mates) has been widely observed to cause aggression in a range of animal species [1]. Aggression is a complex behavior and requires the integration of environmental and internal signals for effective behavioral output [2]. Many environmental factors (*e.g.*, resource value and resource-holding potential (RHP)), genes (*e.g.*, *fruitless* and *Sry*) and neurotransmitters (*e.g.*, serotonin and dopamine) control or influence the onset of aggression, aggressive behavior, and outcomes of aggressive behavior [3–5]. Remarkable advances in the study of aggression have been made in the fields of behavioral ecology, evolutionary biology, and molecular biology, as well as in a wide range of species, from mammals to insects and even nematodes [6–10].

Typically, aggression is regarded as costly in terms of energy consumption, time commitment and increased risk of injury or death. Evolutionarily, selection should favor contestants that weigh the potential benefits and costs of conflict and adjust their behavior accordingly [11]. As conflict is potentially damaging and energetically costly, individuals of most species (including the model species *Drosophila melanogaster*) usually avoid escalation and may give up and retreat before becoming injured. However, few species, mainly belonging to two groups of arthropods (Arachnoidea and Insecta), engage in unusually extreme aggression that results in severe injury or death of contestants [12]. Predictions from “hawk-dove” game theory suggest that extreme forms of aggression are an evolutionarily stable strategy only when critical resources are limited, and the benefits of winning outweigh the potential costs of conflict [3,7]. Additionally, individuals must be equipped with effective weapons for the strategy to be viable (reviewed by Enquist & Leimar 1990) [12]. Studies of extreme aggression have mainly focused on behavioral ecology and evolutionary biology perspectives in a few species of Hemiptera [13], Araneae [14], Hymenoptera [15,16] and Nematoda [17] that appear to be unique and challenge the original predictions, *e.g.*, assessment and adjustment strategies. Animals commonly utilize assessment strategies (*e.g.*, sequential assessment and cumulative assessment models) for gathering information about resources (*e.g.*, value) and opponents (*e.g.*, RHP) and adjust their aggression accordingly [18,19], but these strategies have limited utility and even fail in species that display extreme aggression [20,21].

Parasitoid wasps are insects that are well known for their biological control of pests and that have been employed as important study species (*e.g.*, *Nasonia vitripennis*) for behavioral ecology, molecular biology and evolutionary biology research [6–8,22]. *Anastatus disparis* (Ruschka) (Hymenoptera: Eupelmidae) is a generalist egg parasitoid of several noxious species of Lepidoptera and is considered an important biological control agent of the severe forestry pests *Lymantria dispar* and *Dictyoploca japonica* in China and North America [23]. To acquire mating opportunities, this species frequently exhibits extreme aggression, such as severing their opponent’s feet and antenna with their mouthparts, resulting in severe injury or even death [21,24] (**Figure 1**). Thus, *A. disparis* provides an excellent opportunity and experimental model to study the behavioral ecology and evolutionary biology of extreme aggression.

**Figure 1.**
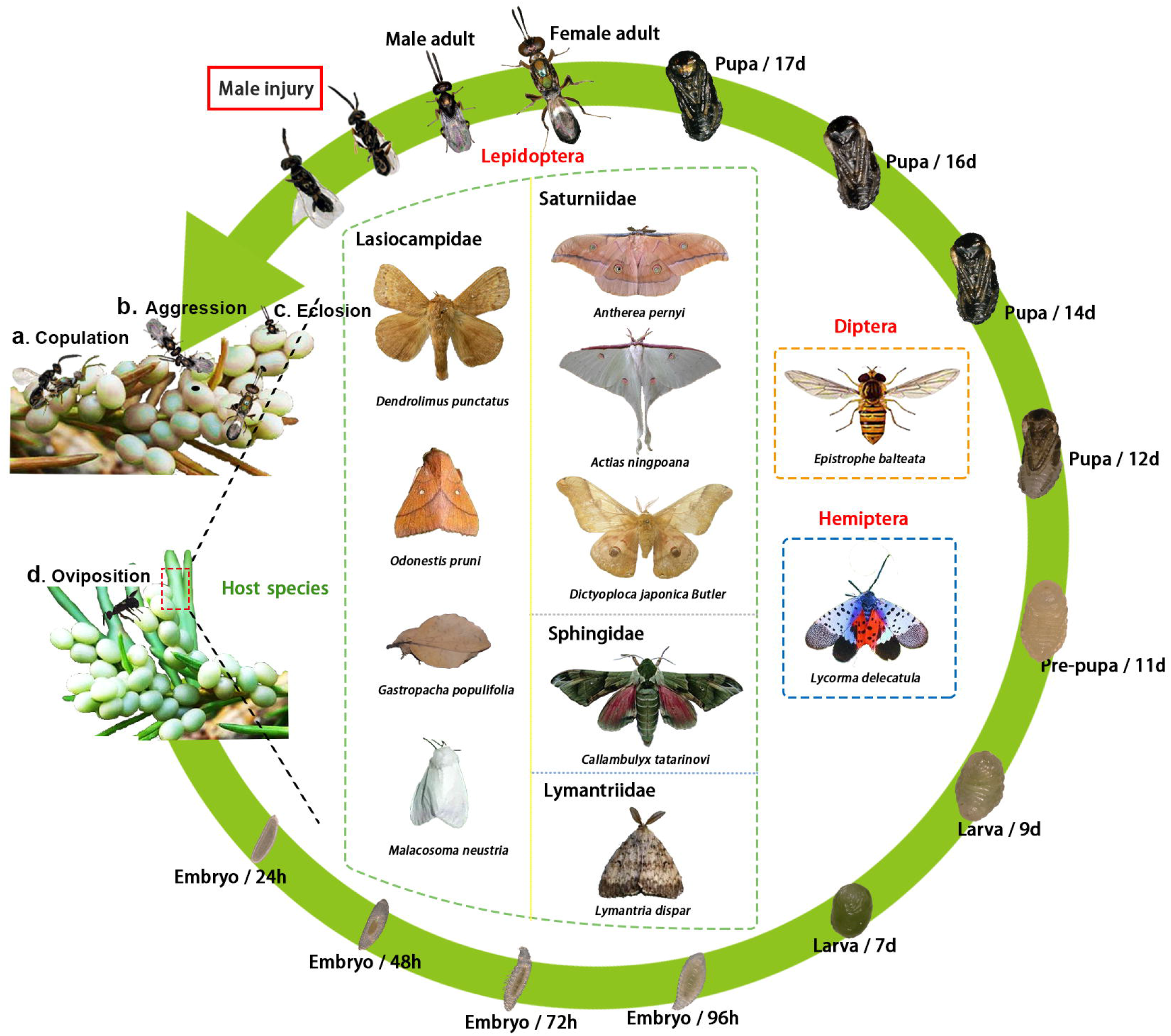
Life cycle and host species of the egg parasitoid wasp *Anastatus disparis*. The developmental period from embryo to adulthood lasts approximately 18 days. Males experience earlier eclosion and remain near the emergence site to find mates and frequently suffer from extreme aggression that results in severe injury or even death. In addition, *A. disparis* has a wide range of host species, including 3 orders and 6 families.

Here, we report a high-quality chromosome-level assembly of the nuclear genome of *A. disparis* that provides comprehensive insight into the molecular background of extreme aggression and the evolutionary basis of such adaptations. First, the basic genome description and comparative genomics of phylogenetic analysis, positively selected genes and gene family expansion were investigated. Aggressive behavior is extremely energetically costly, and such conflicts have been regarded as “energetic wars of attrition” [25,26], as energy expenditure increases as a contest escalates in intensity [27,28]. Thus, integrated with a multiomics analysis of comparative genomics, transcriptomics and behavior, we expected to explore the role of energy metabolism and the selection of energy sources in extreme aggression in *A. disparis.* In general, our study provides insight into molecular and evolutionary studies of extreme aggression and greatly facilitates future analyses of trait evolution in insects. In addition, this here presented genome assembly provides a valuable resource for further research on the molecular basis of evolutionary adaptations such as venom function and behavioral host specificity in *A. disparis.*

## Results

### Genome assembly and annotation

Sequencing of the genome of the extremely aggressive parasitoid wasp *A. disparis* on the Nanopore platform generated 91.36 Gb of clean data with 97.23 × genome coverage after filtering out low-quality sequences and adaptor sequences (Table S1). After error correction of clean reads and assembly polishing, we obtained a high-contiguity genome assembly of 939.58 Mb consisting of 408 contigs with a contig N50 of 5.04 Mb (Table S2). The genome size of *A. disparis* is larger than most of the hymenopteran genomes obtained to date (ranging from 180Mb to 340 Mb) (Table S3) [29]. Consistent with the low GC content reported in hymenopterans (ranging from 30% to 45%) [30], the GC content of the *A. disparis* genome is 29.50%. To improve the genome assembly at the chromosomal level, Hi-C technology was used, and a total of 86 scaffolds representing 99.67% of the total assemblies were successfully anchored to 5 pseudochromosomes (Table S4, Figure S1). The length of the largest chromosome is 220.16 Mb, while that of the smallest is 170.35 Mb (Table S5). The scaffold N50 of the chromosome-level genome assembly reached 183.27 Mb. Based on the both BUSCO result and mapping quality, the *A. disparis* genome assembly displayed high quality and completeness; 99.38% of the *de novo* assembled transcripts were mapped to the reference genome (Table S6). Gene space coverage estimated with the BUSCO pipeline indicated that the percentage of conserved arthropod genes is higher than 96.05% (1,313/1,367) (Table S7).

Currently, the genome sizes reported for most hymenopteran species are moderate, between 180 and 340 Mb [29]. However, the genome size of *A. disparis* was approximately twofold larger, reaching 939.58 Mb. The expansion of repeated sequences is one of the most important factors in genome size in many insects [31,32]. Undoubtedly, the extensive repeated sequences (612.90 Mb, 65.23% of the whole genome, Table S8) in the *A. disparis* genome contributed to its increased size. The percentage of repeated sequences is higher than that in most sequenced hymenopteran insects, e.g., *N. vitripennis* (20.63%) and *Trichogramma pretiosum* (30.30%), and similar to that in *Gonatopus flavifemur* (60.70%) [33]. In contrast to the parasitoid *G. flavifemur*, in which the expansion of transposable elements (TEs) with DNA transposons resulted in the increased number of repeated sequences, the large size of the *A. disparis* genome is mainly due to diverse repeat types of RNA transposons (408.10 Mb, 43.43% of the whole genome).

In total, 19,246 protein-coding genes are predicted in the genome of *A. disparis* based on the *de novo* prediction, homology alignment and RNA-seq transcript assembly (Table S9, Figure S2). We used BUSCO to evaluate the completion of our gene annotation and found that 96.20% of the 1367 genes were completely found (Table S7). The number of protein-coding genes in *A. disparis* was higher than that in most parasitoid wasps (Table S3), but is lower than that in *N. vitripennis* (24,388 predicted genes). The predicted numbers of exons and introns are 104,333 and 85,087, respectively, with an average of 5.42 exons per gene and an average of 4.42 introns per gene (Table S10). A total of 434 noncoding RNAs, including miRNAs (40), rRNAs (114), and tRNAs (280), were identified by *in silico* prediction (Table S11). In addition, 3,589 pseudogenes were predicted in the *A. disparis* genome, with an average length of 2,621 bp (Table S12). Approximately 91.56% (17,621) of the predicted genes are functionally annotated in five public databases: nonredundant protein (Nr), Gene Ontology (GO), TrEMBL, euKaryotic Orthologous Groups (KOG), and KEGG Ortholog (KO) (Table S13). It will provide a comprehensive molecular basis for searching for possible evolution adaptations in *A. disparis.*

### Comparative genomics

To better understand the evolutionary events of the *A. disparis* genome through the analysis of major, we performed a comparative analysis with 11 other insect genomes containing nine Hymenoptera (*e.g.*, *Athalia rosae, Atta cephalotes, Apis mellifera, Macrocentrus cingulum, Ceratosolen solmsi, Trichogramma pretiosum, N. vitripennis, Copidosoma floridanum, Fopius arisanus), D. melanogaster* and the red flour beetle *Tribolium castaneum.* The phylogenetic relationships among *A. disparis* and 11 other insect species (*T. castaneum*, as outgroup) were determined according to a genome-wide set of 1,760 single-copy genes. The phylogenetic analysis by maximum likelihood (PAML) showed that five chalcidoids, *A. disparis, N. vitripennis, C. solmsi, T. pretiosum*, and *C. floridanum*, clustered together and that *A. disparis* had a closer relationship to *N. vitripennis* than to the other species. The estimated divergence times suggested that *A. disparis* diverged from the common ancestor of *A. disparis-N. vitripennis* ~79 Mya (95% CI: 57-103) (**Figure 2**A).

**Figure 2.**
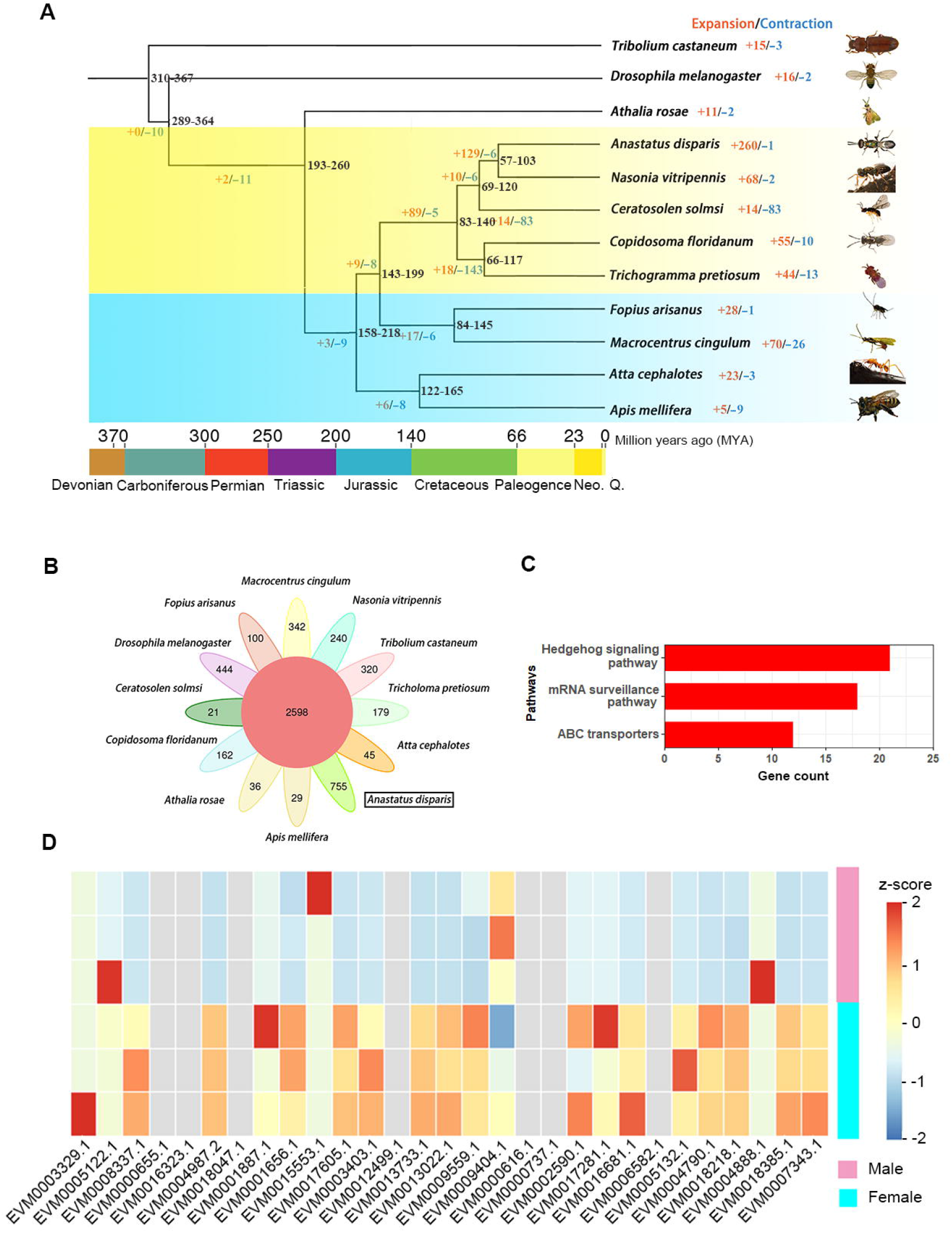
Comparative genomic analysis of the parasitoid wasp *Anastatus disparis*. A. Maximum likelihood tree showing the phylogenetic relationships of *A. disparis*, nine additional species of Hymenoptera, the fly *Drosophila melanogaster* and the beetle *Tribolium castaneum.* The phylogenetic tree was based on 1,760 single-copy proteins, with *T. castaneum* serving as the outgroup. The numbers of expanded (red) and contracted (blue) gene families are shown on the branches. B. The petal figure informs about the number of gene families shared among the twelve investigated species, in which the central circle indicates the number of gene families shared by all species, and the edge circles indicate the number of species-specific gene families. C. Enriched KEGG pathways of *A. disparis*-specific genes. D. Sex differences in the expression of all roadkill genes from the *A. disparis* genome. The enriched Hedgehog signaling pathway contained many species-specific genes, such as roadkill, that display female-biased expression.

We identified a total of 16,159 gene family clusters in our analysis (Table S14). A total of 2,598 gene families are shared by all selected 12 species and 755 *A. disparis-specific* gene families (*i.e.*, lacking 1: 1ortholog in other species) were identified (**Figure 2**B). Interestingly, GO and KEGG enrichment analyses revealed that the *A. disparis*-specific genes were enriched in eye formation (KO: 04341, Hedgehog signaling pathway-fly, *P* < 0.001; GO: 0042067, establishment of ommatidial planar polarity, *P* < 0.001) (**Figure 2**C, Figure S3). Many *A. disparis*-specific genes annotated as roadkill genes regulate the Hedgehog signaling pathway, which participates in the development of numerous tissues and organs [34]. Combined with transcriptomic data of female and male adults (SRA: PRJNA642922), we found that most genes in the Hedgehog signaling pathway were significantly upregulated in adult females, *i.e.*, female-biased genes (Criteria: at least 2-fold expression change of female/male, *P* < 0.05, **Figure 2**D).

### Traces for historical selection in genes

A total of 1,760 single-copy genes were screened for signatures of positive selection on the terminal branch of *A. disparis* in the phylogeny, as shown in Figure 2A. The branch site model in PAML [35] was used to define the significance (*P* < 0.05, false discovery rate (FDR)-adjusted) of each candidate gene. In total, 354 genes showed evidence of positive selection in the *A. disparis* genome. We did not observe any significantly enriched GO terms or KEGG pathways for these positively selected genes (*P* > 0.05). Combined with the transcriptomic analysis (SRA: PRJNA826118) and qRT□PCR analysis, we found several positively selected genes, including *apoLp (apolipophorins*, EVM0014364.1), *LRP4 (low-density lipoprotein receptor-related protein 4*, EVM0015878.1), *PLA2 (phospholipase A2*, EVM0007663.1), *POA3-like (phenoloxidase subunit A3-like*, EVM0007014.1), and *NF-KBIC (NF-kappa-B inhibitor cactus*, EVM0013578.1) in *A. disparis* that were significantly higher expressed in males showing aggressive behavior than in males that were not aggressive.

### Gene family expansions

We used CAFE software (v.4.2) to study gene family expansion and contraction during the evolution of *A. disparis* and related species. By comparing twelve species, we found that the *A. disparis* genome displayed one contracted and 260 expanded gene families compared to the common ancestor of *A. disparis* and *N. vitripennis* (Figure 2A). Enrichment analysis revealed that the expanded genes were mainly associated with detoxification (*e.g.*, drug metabolism-other enzymes (KO: 00983, genes = 23, *P* = 2.46E-11), metabolism of xenobiotics by cytochrome P450 (KO: 00980, genes = 17, *P* = 6.19E-10), and drug metabolism-cytochrome P450 (KO: 00982, genes = 15, *P* = 4.67E-08)) (Figure S4). Cytochrome P450 genes displayed distinct expansion, and 140 cytochrome P450 genes were identified in the *A. disparis* genome, which is greater than the number found in the closely related wasp *N. vitripennis.*

We found several additional gene families that expanded in *A. disparis* since its ancestor split from *N. vitripennis*, e.g., the *4CL1-like (4-coumarate: CoA ligase 1-like*) and *PGRP-SC2-like (Peptidoglycan recognition proteins SC2-like*) families, involved in *A. disparis* aggression (see below transcriptomic analysis of male aggression lasting for 30 mins and 60 mins, SRA: PRJNA826118). Phylogenetic analysis showed that *4CL1-like* genes were largely expanded in the *A. disparis* genome (**Figure 3**A). A total of fifteen *4CL1-like* genes were identified, and the gene family is more diverse in *A. dipsaris* than in any of the other investigated hymenopteran insects in this study. Combined with transcriptomic data of female and male adults (SRA: PRJNA642922), more than half of the *4CL1-like* genes showed sex-biased expression, including seven female-biased genes and one male-specific gene (Criteria: FPKM < 0.4 in females, FPKM > 2 in males [36]). This male-specific gene (EVM0003417.1) showed tissue-specific expression in the male head and was also highly expressed during aggression (Figure 3, LC-MS data of protein from the tissue of male, transcriptomic analysis of male aggression lasting for 30 mins and 60 mins, SRA: PRJNA826118).

**Figure 3.**
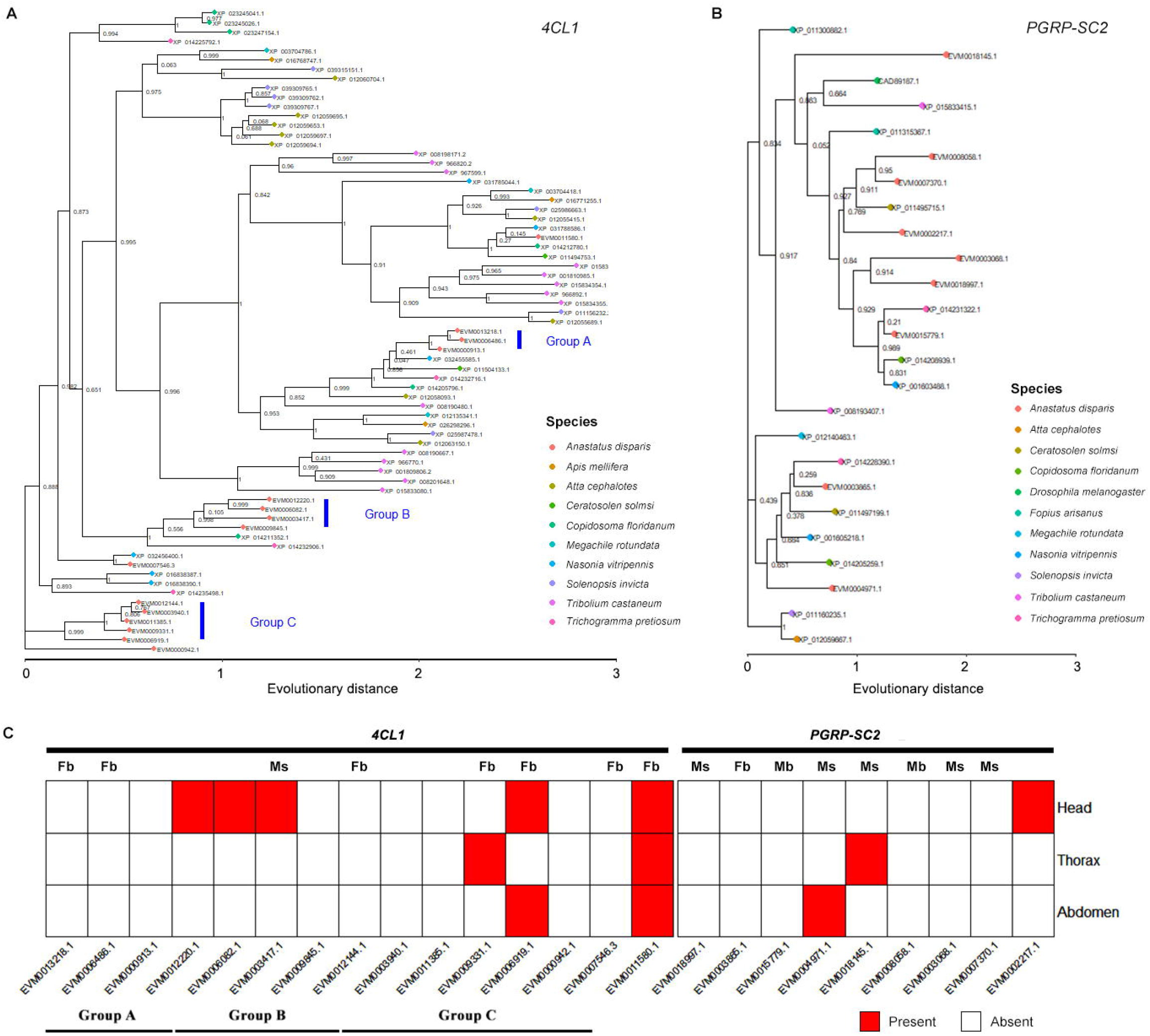
The phylogenetic relationships and expression patterns of expanded *4CL1-like* and *PGRP-SC2-like* genes. A. The maximum likelihood phylogenetic tree of *4CL1-like* proteins in *Anastatus disparis* and other insect species. B. The maximum likelihood phylogenetic tree of *PGRP-SC2-like* proteins in *A. disparis* and other insect species. The maximum-likelihood phylogenetic tree was built using IQ-TREE, and branch support was assessed by 1,000 ultrafast bootstrap replicates. C. The expression pattern of *4CL1-like* and *PGRP-SC2-like* genes in *A. disparis* across sexes and tissues is indicated according to transcriptomic data of female and male adults (SRA: PRJNA642922) and LC-MS data of protein from the tissue of males. Mb/Fb indicates male/female biased genes (FPKM > 1, *P* < 0.05), and Ms/Fs indicates male/female specific genes (FPKM < 0.4 in males and FPKM > 2 in females, FPKM < 0.4 in females and FPKM > 2 in males). FPKM: fragments per kilobase of transcript per million fragments mapped.

*PGRPs* are pattern recognition molecules conserved from insects to mammals that recognize bacteria and their unique cell wall component, peptidoglycan [37,38]. In *D. melanogaster*, 13 genes encoding *PGRPs* have been described and classified on the basis of molecular mass. *PGRP-SC* is one of the short forms with a molecular mass below 20 kDa. In the *A. disparis* genome, we identified nine *PGRP-SC2-like* genes, which are largely expanded and much more abundant than in *Drosophila* species and the closest relative of *A. disparis, N. vitripennis* (**Figure 3**B). Most *PGRP-SC2-like* genes showed male-biased expression (transcriptomic analysis, SRA: PRJNA642922), including one expressed in aggressive males associated with aggression (*i.e.*, EVM0004971.1, transcriptomic analysis of male aggression lasting for 30 mins and 60 mins, SRA: PRJNA826118) (**Figure 3**C).

### Metabolic genes differentially expressed during male-male aggression in the absence of females

Compared to the newly eclosed male without any aggressive experience, 20 and 52 differentially expressed genes (DEGs) were found in males during aggression lasting for 30 mins and 60 mins, respectively (Figure S6&S7, transcriptomic analysis, SRA: PRJNA826118). Some of these genes encode carbohydrate/lipid transporters and metabolic enzymes (**Figure 4**). A *Tret1-like* (trehalose transporter 1-like, EVM0000934.1) gene was significantly upregulated 3-4-fold during the first 30 min of aggressive behavior but was then downregulated in the later aggression period. The cellular membrane is impermeable to trehalose; thus, Tret is a high-capacity trehalose transporter that allows exogenous trehalose uptake into cells [39]. Trehalose is the principal sugar circulating in the blood or hemolymph of most insects and has essential roles in energy storage [40]. In addition to trehalose, other types of sugar, such as glycogen, may also be consumed, and genes encoding *agl-like* (*α-glucosidase-like*, EVM0005860.1), *GP (glycogen phosphorylase*, EVM0014398.1), and *stp-like (sugar transporter-like protein*, EVM0011231.1) are differentially expressed in aggression males. The three main pathways involved in energy metabolism (glycolysis, the tricarboxylic acid cycle, and oxidative phosphorylation) all generate ATP. The glycolysis-related gene *GCDH-like (glucose dehydrogenase [acceptor]-like*, EVM0015375.1) followed a similar pattern and was upregulated in the early stage of aggression and downregulated in the late phase. In addition to carbohydrates, lipids are a major energy source, and the energy from oxidative metabolism is more than twice that produced by the oxidation of an equal weight of carbohydrate [41]. The number of genes associated with steatolysis and transportation, such as *ACBP4 (acyl-CoA-binding domain-containing protein 4*, EVM0016316.1), *4CL4-like, PLA2, iPLA(2) (calcium-independent phospholipase A2*, EVM0004294.2), *Lip1* (Lipase1, EVM0011964.1), and *SCAD (short-chain acyl-CoA dehydrogenase*, EVM0010514.1), were consistently highly expressed from 15 mins to 60 mins, and the expression of the genes *apoLp, Vg3-like (vitellogenin-3-like*, EVM0007464.2), and *LRP4* was upregulated in the later period of aggression. The significant increase in the expression of these genes expedited lipid metabolism from transport to oxygenolysis, producing large amounts of energy for aggression. However, the expression of genes related to fatty acid synthesis, *i.e., FAS-like (fatty acid synthase-like*, EVM0009772.1), *ADΔ11* (*acyl-CoA-Δ(11) desaturase*, EVM0015082.1), and *LPIN1* (*phosphatidate phosphatase LPIN1*, EVM0015404.1), was consistently downregulated in aggressive males (**Figure 4**C). Besides, we found that oxidative phosphorylation (the main pathway involved in energy metabolism)-related genes of *AKR1A1-like (alcohol dehydrogenase [NADP(+)]-like*, EVM0004274.3, EVM0017105.1) were upregulated throughout the period of aggression. These results suggested that, in addition to “carbohydrates”, the energy required to sustain and support aggression in *A. disparis* may also be provided by lipid metabolism.

**Figure 4.**
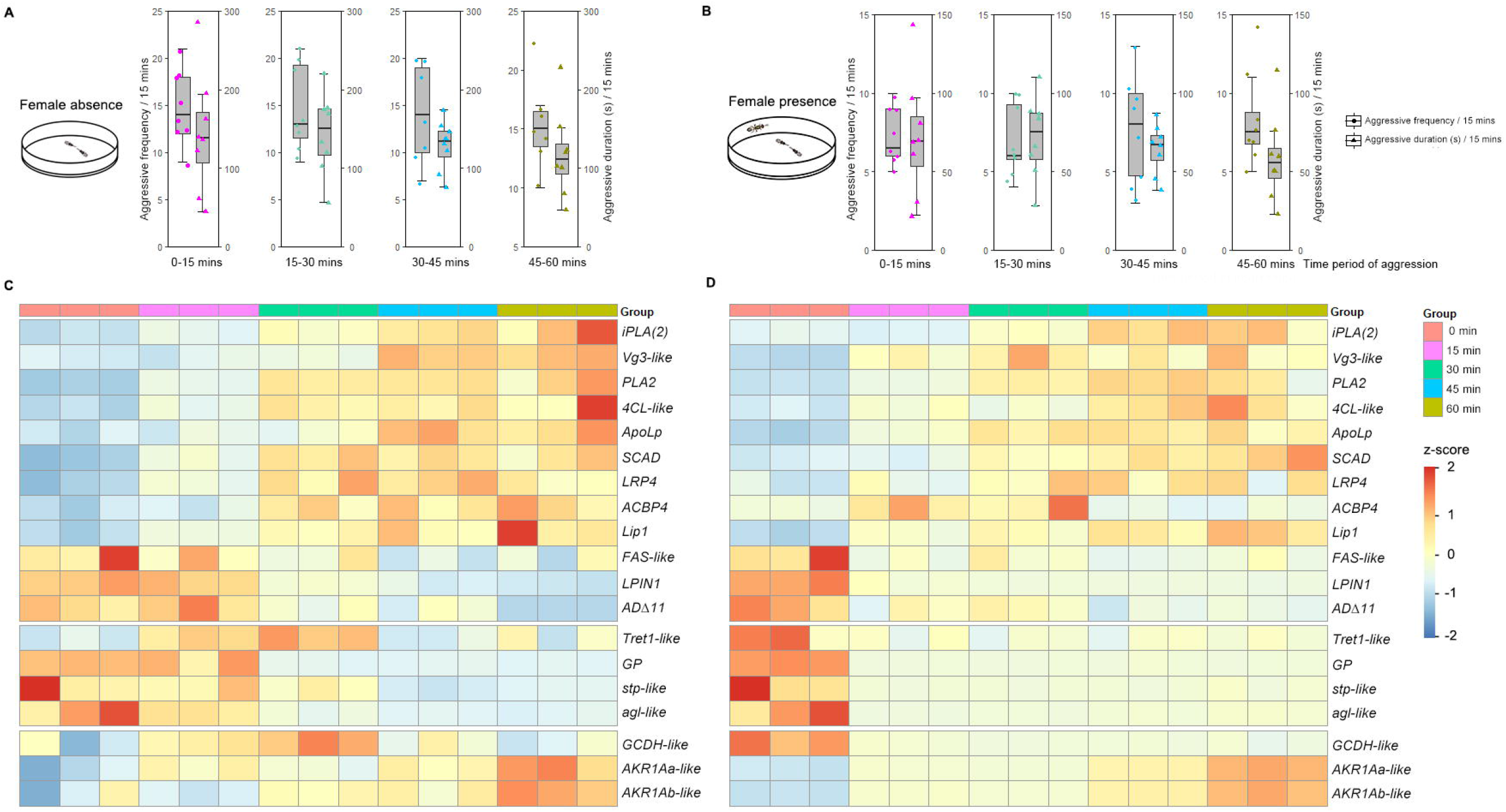
Behavioral aggression and gene expression patterns for metabolism under the conditions of female absence and presence. A. Boxplots describing the frequency and duration of aggression on four consecutive timescales for 60 mins under the condition of female absence. B. Boxplots describing the frequency and duration of aggression on four consecutive timescales for 60 mins under the condition of female presence. C. The expression pattern of metabolic genes at 0, 15, 30, 45 and 60 min under the condition of female absence. D. The expression pattern of metabolic genes at 0, 15, 30, 45 and 60 min under the condition of female presence. Gene expression was determined through qRT-PCR and calculated by the 2^-ΔΔCt^ method using the housekeeping gene EF1A as a reference.

### Changes in metabolic genes related to extreme aggression in the presence of females

When females were present, the frequency (15.53 ± 0.93 per 15 mins) and duration (133.56 ± 9.58 s per 15 mins) of aggressive behaviors increased approximately 3-fold, which was a significant increase compared to aggression in the absence of females (frequency: *F*_(1, 63)_ = 8.205, *P* < 0.001; duration: *F*_(1, 63)_ = 36.831, *P* < 0.001). Similar to the pattern in the absence of females, the frequency and duration of aggression did not significantly change with the passage of time (frequency: *F*_(3, 28)_ = 0.412, *P* = 0.746; duration: *F*_(3, 28)_ = 0.328, *P* = 0.805) (**Figure 4**A & **4**B). The above genes (*PLA2, iPLA(2), 4CL-like, LPIN1, LRP4, apoLp, Vg3-like, Tret1-like*, *FAS-like, AKR1A1-like, ACBP4*, and *GCDH-like*) encoding carbohydrate/lipid transporters and metabolism were also differentially expressed in aggressive males according to qRT-PCR (**Figure 4**D). Although genes related to lipid transport and metabolism were consistently highly expressed until the end of the experiment (*i.e.*, 60 mins), the expression of sugar-related genes was downregulated earlier (*i.e.*, after 15 mins) in the presence of females than in their absence. To support the significant increases in the intensity [21] and frequency of male *A. disparis* aggression in the presence of females (Figure 4), carbohydrates (*e.g.*, trehalose and glycogen) might be quickly consumed, and lipid metabolism might provide more energy during more intense and/or frequent aggression.

### Changes in metabolic genes in winners and older males

This study screened a series of genes associated with metabolism during aggression; these genes are also likely to play an important role in other models of aggression. First, an individual’s RHP determines their potential to win contests based on morphological and physiological traits [42]. Compared to *A. disparis* males that lost contests, several carbohydrate and lipid metabolism genes (*agl-like, Tret1-like, stp-like, apoLp*, and *4CL-like*) were significantly more highly expressed in winners (**Figure 5**A, transcriptomic analysis, SRA: PRJNA826557). Similar to many studies [43], energy metabolism ability is likely an influential RHP component in *A. disparis*, and the winners likely consumed more energy produced by the oxidation of carbohydrates and lipids to win the contests. In losers, the upregulated genes were found to be involved in the immune response to bacterial genes, *e.g., NFkBIC, PPI (pacifastin-like protease inhibitor cvp4*, EVM0008147.1), and *POA3-like.* Extreme aggression in *A. disparis* males typically ends with losers being injured while winner fewer being injured [44]. As the possibility of injury increases for individuals that lose conflicts, injuries from aggression might expose the individual to exogenous pathogens and activate the immune response in this case.

**Figure 5.**
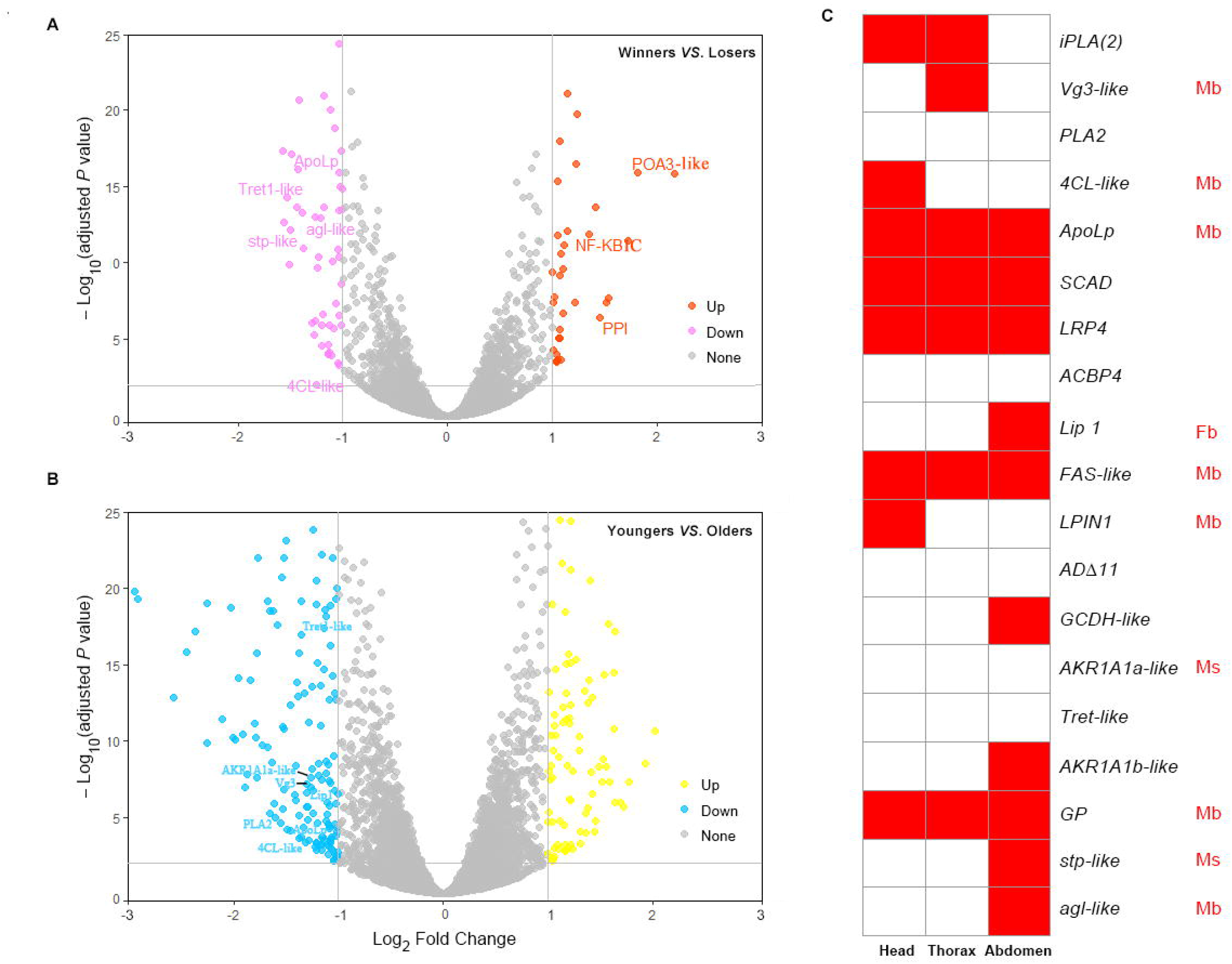
Differentially expressed metabolic genes in winning and older males. A. Volcano plot showing differential gene expression between winners and losers. There were 42 significantly upregulated genes (shown in red) and 59 significantly downregulated genes (shown in pink). Compared to losers, several metabolic genes, including *agl-like, Tret1-like, stp-like, ApoLp*, and *4CL-like*, significantly increased their expression, and that of the antibacterial genes *NFkBIC, PPI*, and *POA3-like* significantly decreased in winners. B. Volcano plot showing differential gene expression between 1-day-old males and 3-day-old males of *A. disparis.* Compared to 3-day-old males, there were 109 significantly upregulated genes (shown in yellow) and 180 significantly downregulated genes (shown in blue) in 1-day-old males. The expression of several metabolic genes, *PLA2, 4CL-like, apoLp, Lip1, AKR1A1-like, Tret1-like* and *Vg3-like*, in 1-day-old males was significantly decreased in 3-day-old males. C. The expression pattern of metabolic genes across different tissues and sexes indicated by transcriptomic data of female and male adults (SRA: PRJNA642922) and LC–MS data of proteins from the tissue of males. Mb/Fb indicates male/female biased genes (FPKM > 1, *P* < 0.05), and Ms/Fs indicates male/female specific genes (FPKM < 0.4 in males and FPKM > 2 in females, FPKM < 0.4 in females and FPKM > 2 in males). FPKM: fragments per kilobase of transcript per million fragments mapped.

Second, aggressive behavior typically has lifetime consequences [45], and empirical reports and theoreticians have revealed that the propensity for aggression tends to increase with age [46]. Younger individuals have higher residual reproductive value than older individuals and have more to lose from injuries than older individuals. However, when *A. disparis* was used as an experimental model to explore the characteristics of aggression from a life-history perspective, the results showed that the frequency and intensity of aggression in *A. disparis* significantly decreased with male age [47]. The genes *PLA2, 4CL-like, apoLp, Lip1, AKR1A1-like, Tret1-like*, and *Vg3-like* were all downregulated in 3-day-old males (lifespan: ~7 days in the wild) (**Figure 5**B, transcriptomic analysis, SRA: PRJNA707227), which might contribute to the unusual life-history aggression pattern of decreasing with male age [47] in this species. In addition, our study also supplies an alternative perspective (that of energy metabolism and physiological state) that might explain the aggression pattern.

### Role of *apoLp* in aggression

According to the above transcriptomic data of males during aggressive behavior (in the condition of female absence and presence), winners and older males, we found a shared DEG, *apoLp*, which was also positively selected and male biased. Phylogenetic analysis showed that this gene was more closely related to that of *N. vitripennis* (**Figure 6**A). The *apoLp* gene encoding apolipophorins is known to play an important role in lipid transport [48,49]. The above findings suggest that energy from the lipid metabolism process likely plays a role in sustaining and supporting intense/frequent extreme aggression in *A. disparis.* Thus, *apoLp* is an essential candidate gene related to aggression in *A. disparis*, and its role was tested as described below.

**Figure 6.**
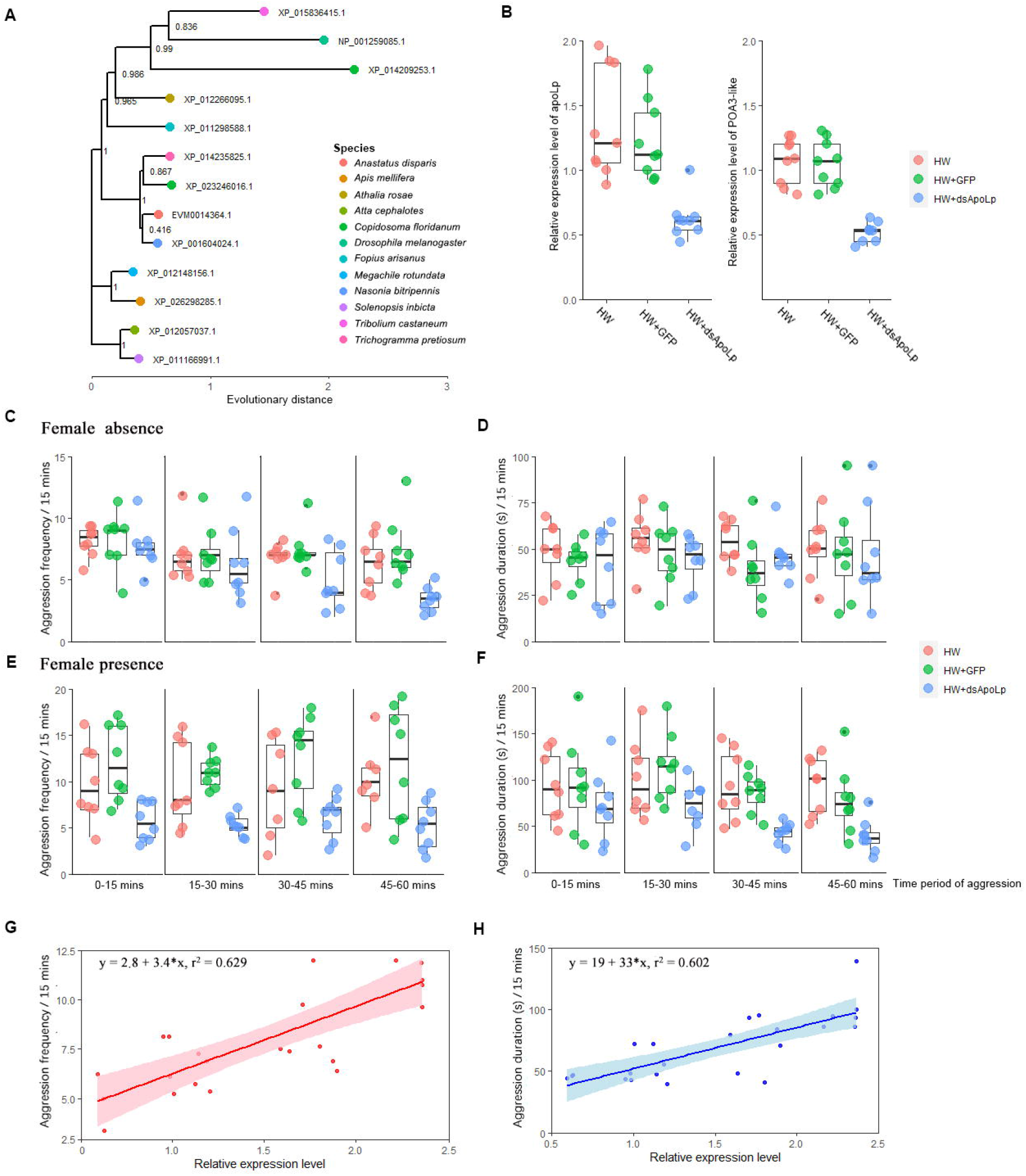
Effect of *apoLp* on aggression. A. The maximum-likelihood phylogenetic tree of *apoLp* proteins in *Anastatus disparis* and eleven other insect species. The tree was constructed using IQ-TREE, and branch support was assessed by 1,000 ultrafast bootstrap replicates. B. The expression of *apoLp* and *POA3-like* genes after males were fed honey water (HW), HW + dsRNA of green fluorescent protein (*dsRNAGFP*) and HW + *dsRNAapoLp.* Gene expression was determined through qRT-PCR and calculated by the 2^-ΔΔCt^ method using the housekeeping gene EF1A as a reference. C. Aggression was described according to the frequency of aggression under the condition of female absence. D. Aggression was described according to the duration of aggression under the condition of female absence. E. Aggression was described according to the frequency of aggression under the condition of female presence. F. Aggression was described according to the duration of aggression under the condition of female presence. G. Pearson’s correlation between frequency of aggression and gene expression of *apoLp.* H. Pearson’s correlation between aggression duration of aggression and gene expression of *apoLp.*

After 3-day-old males were fed double-stranded RNAs (dsRNAs) targeting *apoLp*, the expression of the *apoLp* gene was significantly downregulated by approximately 50% compared to that of males feeding on honey water (HW) (*t* = 4.913, *df* = *16, P* < 0.001) and HW + *dsRNA* of green fluorescent protein (*dsRNAGFP) (t* = 5.378, *df* = 16, *P* < 0.001) (**Figure 6**B). The behavioral results showed that except for the aggression duration in the absence of females (HW *vs*. HW + *dsRNAapoLp: χ* = 0.736, *P* = 0.391; HW + *dsRNAGFP vs.* HW + *dsRNAapoLp: X* = 0.148, *P* = 0.7), the frequency and duration of aggression in males significantly decreased after downregulation of *apoLp* expression in both conditions: the absence (frequency: HW *vs*. HW + *dsRNAapoLp: X* = 5.676, *P* = 0.017; HW + *dsRNAGFP vs.* HW + *dsRNAapoLp: X* = 9.676, *P* = 0.002) and presence of females (frequency: HW *vs*. HW + *dsRNAapoLp: X* = 46.145, *P* < 0.001, HW + *dsRNAGFP vs.* HW + *dsRNAapoLp: X* = 68.734, *P* < 0.001; duration: HW *vs*. HW + *dsRNAapoLp: X* = 10.939, *P*=0.001, HW + *dsRNAGFP vs.* HW + *dsRNAapoLp: X* = 15.069, *P* < 0.001) (**Figure 6**C & **6**D). Furthermore, the frequency of aggression in males decreased after 15 min of feeding on *dsRNAapoLp* (*vs*. after 45 min in the absence of females). In addition, Pearson’s correlation analysis revealed that the gene expression of *apoLp* was moderately positively correlated with aggression frequency (**Figure 6**E) and duration (**Figure 6**F). The above results provide supporting evidence that genes related to lipid metabolism likely play a role in the aggressive behavior of *A. disparis*, especially for late- and high-intensity aggression.

In addition to lipid transport, *apoLp* has been reported to increase antibacterial activity, participate in activation of the prophenoloxidase cascade and participate in cellular immune responses [50,51]. Consistent with these roles, one prophenoloxidase pathway gene, *POA3-like*, was significantly downregulated after males were fed dsRNA-apoLp (HW *vs*. HW + *dsRNAapoLp: t* = 5.741, *df* = 10, *P* < 0.001; HW + *dsRNAGFP vs.* HW + *dsRNAapoLp: t* = 6.27, *df* = *10*, *P* < 0.001) (Figure 6B).

## Discussion

In this study, a high-quality chromosome-level genome assembly of *A. disparis* was provided, and we found that genome size distinctly enlarged approximately twofold (compared to most hymenopterans), reaching 939.58 Mb. The low GC content of 29.50% in the *A. disparis* genome may be caused by GC-biased gene conversion and high recombination rates [52]. Similar to other insect species, the expansion of repeated sequences (mainly TEs with RNA transposons: 408.10 Mb, 43.43% of the whole genome) was one of the most important factors in increasing the size of the *A. disparis* genome. TE expansions and insertions can cause a variety of changes in the host genome and might contribute to adaptation, *e.g.*, helping insects to increase insecticide resistance and adapt to climate change [53,54]. The above environmental risks are frequently encountered by *A. disparis* due to the widespread application of insecticides and the wide distribution of its host species, from temperate to frigid zones [23].

Comparative genomics analysis revealed that many *A. disparis-specific* genes annotated as roadkill genes regulate the Hedgehog signaling pathway. This regulation may be important during eye formation for the proper packaging of ommatidia into a hexagonal array in flies. In humans, a dramatic effect of reduced Hedgehog signaling in embryos is cyclopia [55]. Sexual dimorphism in some wasp species (*e.g.*, *Melittobia* spp.) is pronounced in the eye; males are blind, whereas females have fully functioning eyes [56]. Combined with most roadkill genes being significantly upregulated in adult females, our genome data suggest that *A. disparis* might exhibit distinct eye development or formation characteristics in females.

The cytochrome P450 gene family displayed distinct expansion in the *A. disparis* genome. Cytochrome P450 enzymes can metabolize a broad range of endogenous and exogenous compounds, thus contributing to many processes, including nutrition provision, growth, development, xenobiotic detoxification, and pesticide resistance [57,58]. As the diverse host range of *A. disparis* contains 3 orders and 6 families (Figure 1) [23], the expansion of cytochrome P450 genes in *A. disparis* might have occurred during evolution to deal with different host metabolites [59] and complex living environments. In addition, this expansion might play an important role in the metabolic biotransformation of insecticides [60]. Thus, these potential genes might also be candidates for resistance in *A. disparis* that would improve its utilization for biological control and the evaluation thereof.

We also found two expanded gene families of *4CL1-like* and *PGRP-SC2-like*, involved in *A. disparis* aggression. *4CLs* have been sequenced from numerous plant species and catalyze the activation of 4-coumarate and related substrates to the respective CoA esters; they play a vital role in the phenylpropanoid-derived compound pathway [61]. However, based on the whole genome shotgun sequence, *4CLs* have also been identified in other insects, such as Hymenoptera, Coleoptera and Lepidoptera species [62]. *4CL* belongs to a class I adenylate-forming enzyme superfamily. This group of enzymes is involved in an immense variety of metabolic pathways, such as fatty acid oxidation and enzyme regulation [63]. *4CL1-like* may serve to synthesize a long chain acyl-CoA in the activation step that ultimately leads to degradation of fatty acids [62,64]. One of the male/head-specific *4CL1-like* genes (EVM0003417.1) was highly expressed during aggression, which likely contributes to the timely provision of enough energy to cut off the limbs of opponents with mouthparts during aggressive behavior. The association of *4CL1-like* with an aggression gene (EVM0003417.1) exhibiting sex and tissue specificity may be the result of long-term evolutionary selection that facilitates adaptation for extreme aggression.

Aggressive behaviors are considered extremely energetically costly [25,26], and several carbohydrate and lipid metabolism genes (*e.g.*, *PLA2, iPLA(2), 4CL-like, LPIN1, LRP4, apoLp, Vg3-like, Tret1-like, FAS-like, AKR1A1-like, ACBP4*, and *GCDH-like*) related to aggression were found in *A. disparis.* Among them, genes of *apoLp, PLA2*, and *LRP4* involving in lipid transport and regulation of lipid metabolism were positively selected genes. Energy to sustain and support aggression in *A. disparis* was provided by carbohydrate and lipid metabolism, while varied with the progress and intensity of the aggressive behavior. Carbohydrates (*e.g.*, trehalose and glycogen) in *A. disparis* may be quickly consumed and provide indispensable energy for early aggression (*i.e.*, the first 30 min of aggressive behavior in the absence of females). Lipid metabolism may provide more energy during later periods of aggression and more intense and/or frequent aggression (*e.g.*, male aggression in the presence of females). Furthermore, males decreased aggression after feeding on *dsRNAapoLp* downregulated the expression of the *apoLp* gene related to lipid transport [48,49], providing supporting evidence.

Typically, evolution favors contestants that accurately assess information about the environment and opponents and then carefully select the appropriate behavior (*e.g*., give up and retreat) [3–5]. However, such behaviors do not exhibit in cases of extreme aggression, where contestants do not retreat and can be severely injured or killed. Our genome data, integrated with multiomics, provide new perspectives for understanding the evolution of energy adaptation in species that exhibit extreme aggression. Energy is presumably the limited resource that motivates a large body of evolutionary theories focusing on energetic trade-offs in behavior [65,66]. Aggressive behaviors are extremely energetically costly and have been considered “energetic wars of attrition” [25,26]. Our study provides evidence that a number of genes involved in carbohydrate/lipid oxidative metabolism are significantly differentially expressed during aggression in *A. disparis.* Links between energy metabolism and aggression are well studied, and a high metabolic rate at rest is often positively correlated with aggression [27,28]. Due to the high-intensity and high energy costs of extreme aggression, energy metabolism is likely the major aspect that favored adaptation for extreme aggression and shaped evolution in *A. disparis.* Furthermore, several metabolic genes related to aggression displayed sex- (*i.e.*, male-) and even tissue-specific expression (Figure 5C), which likely reflect other aspects of the adaptation for male-male aggression.

Carbohydrate uptake and metabolism are important sources of energy that affect aggression in many species [26,43,46,67,68]. In contrast, energy for sustaining and supporting extreme aggression in *A. disparis* is more likely provided by lipid oxidative metabolism, probably due to local mate structure. As its hosts are spatially aggregated (*e.g.*, egg masses of *L. dispar* and clutches of *Antheraea pernyi* eggs) [23,69], *A. disparis* is a classic quasi-gregarious species. Similar to most gregarious species (*N. vitripennis* [22]), *A. disparis* females disperse after mating [23], and newly eclosed males aggregate and wait for female eclosion near the emergence site (*i.e.*, local mate structure) [6], during which male-male chasing and aggression occur [21]. In this mate structure, males rarely leave the emergence site [22,70], which leads to a lack of available exogenous carbohydrates, *i.e.*, honeydew. Thus, evolutionary selection for the use of lipids as an important energy source for the adaptation for extreme aggression in *A. disparis* might be caused by the species’ local mate structure.

In contrast to the aggressive behavior of most species, extreme aggression is usually characterized by conflicts that end with severely injured or dead contestants [21,24]. Consequently, injuries from extreme aggression might increase the probability of the individual being exposed to exogenous pathogens. Obviously, antibacterial activity was activated in aggressive males (upregulated male-biased *PGRP-SC2-like* involved in recognizing pathogens [37,38]) and in losers (upregulated genes of *NFkBIC, PPI*, and *POA3-like* involved in the immune response to bacteria). Furthermore, in addition to lipid transport, *apoLp* has been reported to increase the antibacterial activity of the prophenoloxidase cascade [50,51], and *POA3-like* was significantly downregulated after males were fed *dsRNAapoLp.* In addition, the innate immune signaling pathway of prophenoloxidase can also be activated by PGRPs (reviewed by Cytryńska *et al.* 2016) [71]. The prophenoloxidase pathway might be activated through the interaction of *apoLp* and *PGRP*, which remains to be explored in *A. disparis* aggression

Currently, a large body of theoretical work in behavioral ecology has focused on the ways in which energetic trade-offs shape behavior over evolutionary time [65,66,68,72]. Numerous molecular biology of aggression mainly focus on the genes involved in sex determination (*e.g.*, *fruitless* and *Sry*) and neurotransmitters (*e.g.*, serotonin, dopamine, and octopamine) that can control or influence aggressive behavior [4,5]. Our study provides insight into molecular and evolutionary studies of extreme aggression in aspects of energy metabolism and energy sources. In addition, the frequency of injury from extreme aggression is also involved in the adaptation to aggression as it activates antibacterial activity, which has rarely been studied in other aggression studies. Few insect species engage in extreme aggression [13,15,16], and genomes of related species have not been provided. Therefore, a high-quality genome of *A. disparis* can greatly facilitate future analyses of extreme aggression trait evolution in insects. Finally, this genome also provides a valuable genomic resource for further research on the molecular basis of evolutionary adaptations such as venom function and behavioral host specificity in *A. disparis* [29,73].

## Methods

### Parasitoid

An *Anastatus disparis* colony was first established from a population reared on *Lymantria dispar* egg masses collected in March 2012 from Longhua County, Hebei Province, China (41°3′N, 117°74′E), and subsequently maintained on *Antheraea pernyi* egg hosts. During the eclosion period, *A. disparis* adults from the 140^th^ to 145^th^ generations in the laboratory were collected for experiments.

### DNA and RNA extraction

Genomic DNA was extracted from approximately 100 female adults using the cetyltrimethylammonium bromide (CTAB) method. The quality and concentration of the extracted genomic DNA was assessed using 1% agarose gel electrophoresis and a Qubit fluorimeter (Invitrogen, Carlsbad, CA). This high-quality DNA was used for subsequent Nanopore and Illumina sequencing. RNA was extracted with TRIzol Reagent (Catalog No. 15596026, Invitrogen, Carlsbad, CA), and first-strand cDNA was synthesized using a PrimeScript RT Reagent Kit (Catalog No. RR037A, TaKaRa, Tokyo), which was used for subsequent transcriptomic analyses, dsRNA synthesis and qRT-PCR.

### Genome sequencing

A total of 15 μg of extracted genomic DNA was sheared using a Megaruptor (Diagenode, Denville, NJ). The large DNA fragment was recycled by BluePippin (Sage Science, Beverly, MA) and then subjected to DNA repair by NEBNext FFPE DNA Repair Mix (Catalog No. M6630L, NEB, Ipswich, MA) and dA tailing by NEBNext Ultra II End-Repair/dA-tailing Module (Catalog No. E7546, NEB, Ipswich, MA). Then, ligation was performed by adding Adaptor Mix (Catalog No. SQK-LSK109, Oxford Nanopore Technologies, Oxford), and a 1-μl aliquot was quantified with a Qubit 3.0 Fluorometer (Invitrogen, Carlsbad, CA). The purified library was loaded onto flow cells for sequencing on a PromethION (Biomarker, Beijing, China) generating raw data. Base-calling analysis of unprocessed data was conducted using Guppy software (v.3.1.5).

### Genome assembly and scaffolding with Hi-C

Nanopore long reads, with a read N50 of 20,733 and a mean read length of 15,928 bp, were used for initial genome assembly. Error correction of clean data was conducted using Canu (v.1.5) [74], and then assembly was performed with SMARTdenovo software (https://github.com/ruanjue/smartdenovo). The consensus assembly was corrected for three rounds of Racon polishing [75] and three rounds of Pilon polishing [76] using the Illumina reads with default settings.

The Hi-C technique was applied to improve the genome assembly to the chromosomal level. The Hi-C [77] library was prepared using the standard protocol modified for application to whole insects [78]. In brief, 100 female adults were dissected into small pieces and immersed in 2% formaldehyde for crosslinking. Then, isolated nuclei were digested with the restriction enzyme Dpn II. The digested fragments were marked by incubating with biotin and ligated to each other to form chimeric circles. The biotinylated circles were sheared and sequenced using the Illumina HiSeq platform with 150 bp paired-end reads. In total, 623.65 million paired-end reads were generated from the libraries, and low-quality sequences and adaptor sequences were filtered out, generating clean data using fastp [79]. By using HICUP (v.0.8.0), the clean paired-end reads were mapped to the draft assembled sequence [80], and a total of 106.79 million uniquely mapped pair-end reads were generated, of which 74.41% were determined to be valid interaction pairs. Combined with the valid Hi-C data, ALLHIC (v.0.9.8) was applied to anchor the contigs onto the linkage groups using the agglomerative hierarchical clustering method [81]. A total of 532 contigs, representing 99.67% of the total assemblies, were successfully anchored to chromosome-level scaffolds [82]. Finally, the scaffolds were concatenated with 100 Ns to create five pseudochromosomes.

BUSCO (v.4.0) [83] with 1,367 genes in Insecta OrthoDB (v.10) was used to assess the completeness and accuracy of the assembled genome. To further assess its completeness, we employed Illumina RNA-seq data to map the assembled genome using bwa software with the software’s default parameters [84].

### Repeat annotation

Two complementary methods of de novo prediction and homology-based searches were used to predict repetitive sequences within the *A. disparis* genome. A de novo repeat libraries were constructed using RepeatScout (v. 1.0.5) [85] and LTR-FINDER (v.1.07) [86] with default parameters, and these repetitive sequences from the libraries were classified using PASTEClassifier (v.1.0) with default parameters [87]. Then, the *de novo* repeat libraries were combined with the Repbase database to obtain the final repeat database [88]. Finally, RepeatMasker (v.4.0.5) with the parameters -nolow -no_is -norna -engine wublast was used to identify repeat sequences in the *A. disparis* genome by aligning them against the final repeat database [89].

### Protein-coding gene, noncoding RNA and pseudogene prediction

For the prediction of protein-coding genes in the *A. disparis* genome, three methods of de novo prediction, homology-based search, and RNA-seq-based assembly were applied. Specifically, the GENSCAN [90], Augustus (v.2.4) [91], GlimmerHMM (v.3.0.4) [92], GeneID (v.1.4) [93], and SNAP (v.2006-07-28) [94] software programs were used for *de novo* prediction with the software’s default parameters. For homology prediction, GeMoMa (v. 1.3.1) [95] software was conducted with amino acid sequences from *Apis mellifera, Athalia rosae, N. vitripennis* and *Macrocentrus cingulum.* For the RNA-seq-based prediction, RNA-seq data from female and male *A. disparis* adults (three replicates for each sex, SRA: PRJNA642922) were mapped to the *A. disparis* genome using HISAT2 (v.2.0.4) [96] and assembled by StringTie (v.2.0) [97], and GeneMarkS-T (v.5.1) was used to predict genes based on the assembled transcripts. In addition, gene prediction based on unigenes assembled by Trinity (v.2.11) [98] was performed by PASA (v.2.0.2) [99]. Finally, all gene models were integrated using EVM (v. 1.1.1) to obtain a consensus gene set [100]. We evaluated the completeness of this final set of protein-coding genes with BUSCO (v.4.0), applying the protein mode by setting “-m proteins”.

MicroRNAs and rRNAs in the assembled *A. disparis* genome were identified by BLASTN searches against the Rfam database (v.12.0) [101]. tRNAs were predicted using tRNAscan-SE (v.1.3.1) with eukaryote parameters [102]. For pseudogene prediction, candidate homologous amino acid sequences in the genome were identified by GenBlastA (v. 1.0.4) with the parameter -e 1e-5 to mask protein-coding genes [103]. Then, pseudogenes with a premature stop and/or frameshift mutation in the coding region were predicted by GeneWise (v.2.4.1) with default parameters [104].

Gene functions were assigned through BLAST searches based on the best match of the alignments against multiple functional databases of GO, KEGG, KOG, TrEMBL, and Nr databases.

### Comparative genomic analyses

Comparative genomic analysis was performed to analyze intraspecific gene duplication, explore the phylogenetic relationships of different species, and identify species-specific genes. The analysis was based on the twelve following genomes: *A. disparis, Athalia rosae* (http://www.insect-genome.com/data/genome_download/), *Atta cephalotes* (https://www.ncbi.nlm.nih.gov/genome/?term=Atta+cephalotes), *Apis mellifera* (https://www.ncbi.nlm.nih.gov/genome/48?genome_assembly_id=403979), *Macrocentrus cingulum* (http://www.insect-genome.com/data/genome_download/), *Ceratosolen solmsi* (https://www.ncbi.nlm.nih.gov/genome/?term=Ceratosolen+solmsi), *Tribolium castaneum* (https://www.ncbi.nlm.nih.gov/genome/?term=txid7070[orgn]), *Drosophila melanogaster* (ftp://ftp.ensembl.org/pub/release-84/fasta/drosophila_melanogaster/dna/), *Trichogramma pretiosum* (ftp://ftp.ncbi.nlm.nih.gov/genomes/all/GCF/000/599/845/GCF_000599845.2_Tpre_2.0), *Nasonia vitripennis* (https://www.ncbi.nlm.nih.gov/genome/?term=Nasonia+vitripennis), *Copidosoma floridanum* (https://www.ncbi.nlm.nih.gov/genome/?term=Copidosoma+floridanum), and *Fopius arisanus* (https://www.ncbi.nlm.nih.gov/genome/?term=Fopius+arisanus).

Clusters of orthologous genes were classified using OrthoFinder (v.2.4) (diamond, e=0.001) [105] and annotated in the PANTHER database (v.15) [106]. As a result, 16,159 gene families were constructed, and 1760 genes were identified as single-copy orthologous genes for phylogenomic analysis and positive selection analyses. For phylogenomic analysis, the amino acid sequences of each single-copy gene family were independently aligned by MAFFT (v.7.205) with the parameter --localpair --maxiterate 1000, filtered by Gblocks (v.0.91b) with the parameter -b5=h, and then concatenated into one supersequence. Then, a species-level phylogenetic tree was constructed by maximum likelihood (ML) using IQ-TREE (v.1.6.11) [107] with the best model (LG + I + G) estimated by modelfinder to evaluate evolutionary relationships. The MCMCTree (v.1.1) program of the PAML package was used to resolve the divergence times among species [108]. Two calibration divergence time points based on fossil records from TimeTree (http://www.timetree.org/) (*A. mellifera vs. A. cephalotes:* 127-192, *T. castaneum vs. A. cephalotes:* 308-366) were used for divergence time calibration in our phylogenetic tree construction.

Positive selection signals were also detected on the *A. disparis* branch using PAML packages (v.4.9) with optimized branch-site mode. A likelihood ratio test was conducted to compare the alternative model (sites under positive selection along the foreground branch) and the null model (sites under purifying and neutrally selection). We calculated the *P* values by the chi-square test with the FDR, and genes with a *P*-adjusted value < 0.05 were identified as positively selected genes.

Gene family expansion and contraction were studied with CAFÉ software (v.4.2). For each branch and node of the phylogenetic tree, an expanded and contracted gene family with both a family-wide *P* value and a viterbi *P* value ≤ 0.05 was defined as a significantly expanded and contracted gene family [109]. KEGG enrichment analysis and annotation were performed using clusterProfile (v.3.14.0) [110]. To identify expanded Cytochrome P450, *4CL1-like* and *PGRP-SC2-like* gene families, protein sequences of well-annotated insects retrieved from UniProt were used as queries to search against the predicted protein sequences from *A. disparis* and selected insects using BLASTP (v.2.8.1, -evalue 1e-5). The putative cytochrome *P450* and *4CL1-like* proteins were further checked for the presence of their characteristic domains of PF00067 and PF00501 from Pfam. Candidate *PGRP-SC2-like* proteins were checked for the alignments of the N-acetylmuramoyl-L-alanine amidase domain using hidden Markov models (HMM search) [111,112]. Phylogenetic analysis of the assigned genes was performed using maximum likelihood methods with the best model estimated by modelfinder in IQ-TREE (v.1.6.11) [107]. Statistical support for all phylogenetic trees was assessed by ultrafast bootstrap analysis using 1,000 replicates [113].

### Behavioral test

Aggressions in two conditions (the absence and presence of females) were assessed in this study. In each arena (height: 1 cm, diameter: 5 cm), two males (and one female in the female presence condition) were simultaneously placed into the arena for 1 hour. There were eight replicates for each treatment. The entire process was recorded using a video camera. Except where noted, both tested females and males were virgin and one-day-old. Aggression in *A. disparis* mainly included sneak attacks by the opponent when courting a female, boxing and chasing. The number of times and duration the males engaged in aggressive behavior were recorded in each arena. To acquire a sample of aggressive males for transcriptomic analyses, aggression lasting for both 30 and 60 mins was conducted. Additionally, aggression lasting for a series of time points of 15, 30, 45 and 60 min was conducted to acquire samples of aggressive males to evaluate metabolic gene expression during aggression.

### Transcriptomic analyses

In our study, four groups (group one: adults of each sex; group two: males from three samples without any aggressive experience, aggression lasting for 30 mins and 60 mins; group three: males of loser and winner during aggression; group four: one- and three-day-old males) were subjected to transcriptomic experiments. Each sample contained 15-20 virgin adults and three replicates, and all adults were one day old except for group four. For the transcriptomic experiment, 3 μg of total RNA from each sample was converted into cDNA using the TruSeq Stranded mRNA LT Sample Prep Kit (Catalog No. RS-122-2101, Illumina, San Diego, CA) to construct cDNA libraries. To obtain raw reads, cDNA libraries were sequenced on an Illumina HiSeq X Ten platform, and 150 bp paired-end reads were generated. Then, reads containing adapter, poly-N reads and low-quality reads were removed from the raw data by FASTX-Toolkit (http://hannonlab.cshl.edu/fastx_toolkit/), yielding clean reads. All the clean reads were mapped to the *A. disparis* genome using HISAT2 (v.2.0.4) [96]. The FPKM (fragments per kilobase of transcript per million fragments mapped) of each gene was calculated using StringTie (v.2.0) [97]. Genes with at least 2-fold expression changes and *P* (FDR-adjusted) < 0.05 as found by R package DESeq2 (v.1.22.1) using the Wald test were considered differentially expressed. The GOseq R package [114] and KOBAS software (v.2.0) [115] were used to determine the significant enrichment of DEGs in the GO subcategories and KEGG pathways, and an adjusted Q-value < 0.05 was chosen as the significance cutoff.

### Effects of *apoLp* on aggression

dsRNA of *apoLp* was synthesized to study the function of gene during aggression. The nucleotide sequence of *apoLp* was obtained from the transcriptomic data of males during aggression (SRA: PRJNA826118), in which forward and reverse primers with the T7 promoter (5’-TAATACGACTCACTATAGGG-3’) were added at the 5’ ends (Table S15). Using Platinum II Taq Hot-Start DNA Polymerase (Catalog No. 14966025, Thermo Fisher Scientific, Waltham, MA), the PCR products for *apoLp* and green fluorescent protein (GFP) were amplified from cDNA in a Mastercycler Nexus instrument (Eppendorf, Hamburg) with a 2-min initial denaturation at 94 °C followed by 30 cycles of 15 sec at 94 °C, 30 sec at 60 °C, 68 °C for 30 sec, and a final hold at 4 °C. DNA fragments of expected size were stained with SYBR Safe DNA gel stain (Thermo Fisher Scientific, Waltham, MA) on an agarose gel and were subsequently extracted using a Universal DNA purification Kit (Catalog No. DP214-02, TIANGEN, Beijing, China) and then used as a template to synthesize dsRNA with a MEGAscript RNAi Kit (Catalog No. AM1626, Thermo Fisher Scientific, Waltham, MA) following the manufacturer’s protocol. The integrity and concentrations of the synthesized dsRNA were determined by 1.5% agarose/Tris-acetate-EDTA gel and Nanodrop (Thermo Scientific Nanodrop 2000, Waltham, MA). RNA interference was done by feeding newly emerged males 10 ng/μL *dsRNAapoLp* or 10 ng/μL *dsRNAGFP* dissolved in 30% honey water for 2 days. Then, the 3-day-old males were divided into two groups to conduct behavioral experiments (n = 8 for each treatment) and qRT-PCR (n = 9 for each treatment).

### qRT-PCR analysis

A series of metabolic gene expression during aggression (n = 3 for each treatment) and gene expressions of *apoLp* and POA3-like after male feeding on *dsRNAapoLp* were evaluated by qRT-PCR analysis (ABI StepOne Plus, Foster, CA) using TB Green Premix Ex Taq II (Catalog No. RR820A, Takara, Tokyo). Relative gene expression was calculated by the 2^-ΔΔCt^ method using the housekeeping gene translation elongation factor 1-α (*EF1A*) as a reference to eliminate sample-to-sample variations in the initial cDNA samples. Primers for the genes were designed using Primer Express 2.0 software (Applied Biosystem, Foster City, CA) (Table S16).

### LC-MS analysis

Protein from the tissue of male heads, thoraxes, and abdomens (approximately 200 adults of one-day-old virgin males were dissected) was extracted by a Total Protein Extraction Kit for Insects (Catalog No. SD-001/SN-002, Invent Biotechnologies, Plymouth, MN, Figure S8). Then, protein was digested with Trypsin Gold (Catalog No. PRV5280, Promega, Madison, WI) at a 1:50 enzyme-to-substrate ratio for 16 hours at 37 □. After desalting and drying, proteomics analyses were performed using an Ultimate 3000nano HPLC system (Thermo Fisher Scientific, Waltham, MA) coupled with a Q-Exactive plus mass spectrometer (Thermo Fisher Scientific, Waltham, MA) operating in data-dependent acquisition (DDA) mode (Figure S9). Finally, using Proteome Discover (v.2.5) and Mascot (v.2.0) software, proteins were identified by searching against the *A. disparis* proteome based on de novo gene perdition with a > 95% confidence interval.

### Statistical analysis

All analyses were performed with *R* software (v.2.14.1, R Development Core Team). The qRT-PCR data comparing gene expression were analyzed with an independent sample *t* test. Data on aggression frequency and duration were analyzed with generalized linear models (GLMs) using the lme4 package. Frequency data were analyzed using a Poisson distribution (quasi-Poisson distribution for overdispersion) and log link function. Duration data were analyzed using a GLM with a normal error structure (and log transformation to normalize the residuals if necessary) [116]. The positive/negative relationship between aggression frequency/duration and the expression level of *apoLp* was assessed with Pearson’s correlation analysis.

## Data availability

The raw genome sequencing data have been deposited in the NCBI Sequence Read Archive (SRA: PRJNA693567). The final genome assembly was submitted to NCBI whole genome shotgun (WGS: JAFFSR000000000). The transcriptome data are available in NCBI SRA (SRA: PRJNA642922, PRJNA707227, PRJNA826118 and PRJNA826557).

## CRediT authorship contribution statement

**Pengcheng Liu**: Supervision, Funding acquisition, Writing - original draft, Writing - review & editing. **Ziyin Wang**: Investigation, Formal analysis, Writing - original draft. **Yumei Tao**: Investigation, Data curation. **Siyu Yin**: Investigation, Data curation. **Jianrong Wei**: Resources, Writing - review & editing. **Jianjun Wang**: Writing - review & editing. **Haoyuan Hu**: Supervision, Writing - review & editing.

## Acknowledgments

This work was supported by the National Natural Science Foundation of China (Grant Nos. 32101534 and 31870639), the Natural Science Foundation of Anhui Province (Grant No. 2108085QC91), Natural Science Foundation of Liaoning Province (2021-MS-053).

## Supplementary material

### Supplementary table titles

**Table S1** Clean data information of *Anastatus disparis* genome.

**Table S2** Primary genome assembly of *Anastatus disparis.*

**Table S3** Genomic features of selected Hymenopteran insects.

**Table S4** Genome assembly at the chromosomal level by Hi-C.

**Table S5** Length of Chromosome in *Anastatus disparis.*

**Table S6** Assessment the gene coverage with the transcriptome data.

**Table S7** Quality assessment of genome assembly and annotation using BUSCO.

**Table S8** Repeat sequences in *Anastatus disparis* genome.

**Table S9** Protein-coding genes predicted in the genome of *Anastatus disparis.*

**Table S10** Basic information of *Anastatus disparis* genome.

**Table S11** Noncoding RNAs in *Anastatus disparis* genome.

**Table S12** Pseudogene in *Anastatus disparis* genome.

**Table S13** Functional annotation of *Anastatus disparis* genome.

**Table S14** Gene family clusters among *Anastatus disparis* and 11 other insect species.

**Table S15** Primer sequences for dsRNA synthesis.

**Table S16** Primer pairs used for expression analysis using qRT-PCR.

### Supplementary figure legends

**Figure S1** The genome-wide Hi-C interaction maps of 5 pseudo-chromosomal linkage groups in *Anastatus disparis.*

**Figure S2** Venn diagram of predicted genes based on the three approaches of *de novo* prediction, homology alignment and RNA-seq transcript assembly in the genome of *Anastatus disparis.*

**Figure S3** Enriched GO category of *Anastatus disparis*-specific genes.

**Figure S4** Enriched KEGG pathways of expanded gene families in *Anastatus disparis.*

**Figure S5** Enriched GO category of expanded gene families in *Anastatus disparis.*

**Figure S6** Volcano plot showing differential gene expression between males during aggression lasting for 30 mins and newly eclosed male.

**Figure S7** Volcano plot showing differential gene expression between males during aggression lasting for 60 mins and newly eclosed male.

**Figure S8** Electrophoretogram of proteins extracted from the tissue of male heads, thoraxes, and abdomens.

**Figure S9** Total ions chromatogram (TIC) of proteins from male heads, thoraxes, and abdomens by LC-MS analysis.

