## Supplementary Table 1 for "The Genome of the Wasp *Anastatus disparis* Reveals Energy Metabolism Adaptations for Extreme Aggression"

**Supplementary Table 1. Clean data information of *Anastatus disparis* genome**

| Data type | Sequence Number | Sum Base (bp) | N50 length (bp) | N90 Length (bp) | | Mean Length (bp) | Max Length (bp) | Mean Quality score |
| --- | --- | --- | --- | --- | --- | --- | --- | --- |
| Clean data | 5,735,295 | 91,357,227,837 | 20,733 | | 9,162 | 15,928 | 340,428 | 8.94 |
