## Supplementary Table 2 for "The Genome of the Wasp *Anastatus disparis* Reveals Energy Metabolism Adaptations for Extreme Aggression"

| **Supplementary Table 2. Primary genome assembly of *Anastatus disparis*** | | | | | | |
| --- | --- | --- | --- | --- | --- | --- |
| Contig number (>1Kb) | Contig length (bp) (>1Kb) | Contig N50 (bp) | Contig N90 (bp) | Contig max (bp) | GC content (%) | Gap total length (bp) |
| 408 | 939581021 | 5041440 | 1273406 | 38756149 | 29.50 | 0 |
