## Supplementary Table 3 for "The Genome of the Wasp *Anastatus disparis* Reveals Energy Metabolism Adaptations for Extreme Aggression"

**Supplementary Table 3. Genomic features of selected Hymenopteran insects**

| Genomic Features | *Anastatus*  *disparis* | *Nasonia*  *vitripennis* | *Trichogramma*  *pertiosum* | *Copidosoma*  *floridanum* | *Pteromalus puparum* |
| --- | --- | --- | --- | --- | --- |
| Genome size (Mb) | 939.58 | 295.78 | 195.09 | 553.96 | 338.1 |
| GC content (%) | 29.50 | 41.7 | 39.9 | 36 | 40.62 |
| Repeat content (%) | 65.23 | 20.63 | 30.3 | 36.85 | 40.11 |
| Number of protein-coding genes | 19,246 | 24,388 | 12,928 | 12,143 | 14,946 |
