## Supplementary Table 4 for "The Genome of the Wasp *Anastatus disparis* Reveals Energy Metabolism Adaptations for Extreme Aggression"

**Supplementary Table 4. Genome assembly at the chromosomal level by Hi-C**

| Item | Value |
| --- | --- |
| Total length (bp) | 939,626,903 |
| Total length without N (bp) | 939,582,303 |
| Total number of scaffold | 86 |
| GC content (%) | 29.50 |
| N50 (bp) | 183,274,343 |
| N90 (bp) | 170,354,989 |
| Average (bp) | 10,925,894.22 |
| Median (bp) | 25,000.00 |
| Min (bp) | 633 |
| Max (bp) | 220,163,934 |
