## Supplementary Table 5 for "The Genome of the Wasp *Anastatus disparis* Reveals Energy Metabolism Adaptations for Extreme Aggression"

**Supplementary Table 5. Length of Chromosome in *Anastatus disparis***

| Chromosome | Length(bp) | contig number |
| --- | --- | --- |
| chr1 | 220,163,934 | 105 |
| chr2 | 190,894,825 | 73 |
| chr3 | 183,274,343 | 128 |
| chr4 | 171,821,451 | 71 |
| chr5 | 170,354,989 | 74 |
| chrUnn | 3,117,361 | 81 |
