## Supplementary Table 6 for "The Genome of the Wasp *Anastatus disparis* Reveals Energy Metabolism Adaptations for Extreme Aggression"

|  | **Supplementary Table 6. Assessment the gene coverage with the transcriptome data** | | | | | |
| --- | --- | --- | --- | --- | --- | --- |
| Species | | Total reads | Mapped reads | Mapped (%) | Properly mapped reads | Properly mapped (%) |
| *Anastatus disparis* | | 284,827,588 | 283,068,693 | 99.38 | 276,122,056 | 97.27 |
