## Supplementary Table 7 for "The Genome of the Wasp *Anastatus disparis* Reveals Energy Metabolism Adaptations for Extreme Aggression"

|  | **Supplementary Table 7. Quality assessment of genome assembly and annotation using BUSCO** | | | | | |
| --- | --- | --- | --- | --- | --- | --- |
|  | Complete BUSCOs | Complete and single copy BUSCOs | Complete and duplicated BUSCOs | Fragmented BUSCOs | Missing BUSCOs | Total Lineage BUSCOs |
| Assembly | 1313 (96.05%) | 1291 (94.44%) | 22 (1.61%) | 18 (1.32%) | 36 (2.63%) | 1,367 |
| Annotation | 1315  (96.20%) | 1285(94.00%) | 30 (2.19%) | 20 (1.46%) | 32 (2.34%) | 1367 |
