## Supplementary Table 8 for "The Genome of the Wasp *Anastatus disparis* Reveals Energy Metabolism Adaptations for Extreme Aggression"

| **Supplementary Table 8. Repeat sequences in *Anastatus disparis* genome** | | | |
| --- | --- | --- | --- |
| Type | Number | Length | Rate (%) |
| ClassI | 1,029,103 | 408,104,751 | 43.43 |
| ClassI/DIRS | 10,065 | 4,191,484 | 0.45 |
| ClassI/LARD | 430,340 | 142,307,365 | 15.15 |
| ClassI/LINE | 135,492 | 66,459,843 | 7.07 |
| ClassI/LTR/Copia | 89,035 | 40,183,616 | 4.28 |
| ClassI/LTR/Gypsy | 266,574 | 147,199,206 | 15.67 |
| ClassI/LTR/Unknown | 43,635 | 21,629,272 | 2.30 |
| ClassI/PLE | 35,809 | 15,708,255 | 1.67 |
| ClassI/SINE | 8,265 | 2,165,276 | 0.23 |
| ClassI/TRIM | 7,244 | 6,724,265 | 0.72 |
| ClassI/Unknown | 2,644 | 843,246 | 0.09 |
| ClassII | 372,692 | 170,866,284 | 18.19 |
| ClassII/Crypton | 903 | 184,185 | 0.02 |
| ClassII/Helitron | 105,247 | 49,050,157 | 5.22 |
| ClassII/MITE | 15,382 | 6,224,218 | 0.66 |
| ClassII/Maverick | 48,634 | 48,733,715 | 5.19 |
| ClassII/TIR | 161,291 | 57,352,813 | 6.10 |
| ClassII/Unknown | 41,235 | 14,412,330 | 1.53 |
| Protential Host Gene | 33,114 | 9,666,372 | 1.03 |
| SSR | 31,052 | 18,387,924 | 1.96 |
| Unknown | 231,755 | 82,571,895 | 8.79 |
| Total | 1,697,716 | 612,899,621 | 65.23 |
