## Supplementary Table 9 for "The Genome of the Wasp *Anastatus disparis* Reveals Energy Metabolism Adaptations for Extreme Aggression"

| **Supplementary Table 9. Protein-coding genes predicted in the genome of *Anastatus disparis*** | | | |
| --- | --- | --- | --- |
| Method | Software | Species | Gene number |
| Ab initio | Genscan | - | 14,454 |
|  | Augustus | - | 23,159 |
|  | GlimmerHMM | - | 87,660 |
|  | GeneID | - | 16,062 |
|  | SNAP | - | 115,351 |
| Homology-based | GeMoMa | *Apis mellifera* | 9,884 |
|  |  | *Athalia rosae* | 10,982 |
|  |  | *Nasonia vitripennis* | 17,814 |
|  |  | *Macrocentrus cingulum* | 16,530 |
| RNAseq | TransDecoder | - | 16,670 |
|  | GeneMarkS-T | - | 3,071 |
|  | PASA | - | 13,520 |
| Integration | EVM | - | 19,246 |
