## Supplementary Table 10 for "The Genome of the Wasp *Anastatus disparis* Reveals Energy Metabolism Adaptations for Extreme Aggression"

| **Supplementary Table 10. Basic information of *Anastatus disparis* genome** | | | | | | | | | | |
| --- | --- | --- | --- | --- | --- | --- | --- | --- | --- | --- |
| Gene Number | Gene  length | Average Gene length | Exon Length | Average Exon Length | Exon Number | Average Exon Number | CDS  Length | Average CDS length | CDS Number | Average  CDS Number |
| 19,246 | 135,723,588 | 7,052.04 | 38,047,394 | 1,976.90 | 104,333 | 5.42 | 27,512,232 | 1,429.50 | 101,147 | 5.26 |
