## Supplementary Table 12 for "The Genome of the Wasp *Anastatus disparis* Reveals Energy Metabolism Adaptations for Extreme Aggression"

| **Supplementary Table 12. Pseudogene in *Anastatus disparis* genome** | | | |
| --- | --- | --- | --- |
| Software | Number | Total length | Average length |
| GeneWise | 3,589 | 9,408,007 | 2,621.34 |
