## Supplementary Table 13 for "The Genome of the Wasp *Anastatus disparis* Reveals Energy Metabolism Adaptations for Extreme Aggression"

| **Supplementary Table 13. Functional annotation of *Anastatus disparis* genome** | | |
| --- | --- | --- |
| Annotation database | Annotated number | Percentage (%) |
| GO Annotation | 6,502 | 33.78 |
| KEGG Annotation | 6,467 | 33.60 |
| KOG Annotation | 10,003 | 51.97 |
| TrEMBL Annotation | 17,508 | 90.97 |
| Nr Annotation | 17,383 | 90.32 |
| All Annotated | 17,621 | 91.56 |
