## Supplementary Table 14 for "The Genome of the Wasp *Anastatus disparis* Reveals Energy Metabolism Adaptations for Extreme Aggression"

**Supplementary Table 14. Gene family clusters among *A. disparis* and 11 other insect species**

| Item | Acep | Adis | Amel | Aros | Cflo | Csol | Dmel | Fari | Mcin | Nvit | Tcas | Tper |
| --- | --- | --- | --- | --- | --- | --- | --- | --- | --- | --- | --- | --- |
| Number of genes | 10,251 | 19,246 | 9,881 | 10,035 | 12,015 | 9,702 | 13,576 | 10,870 | 11,993 | 13,578 | 12,814 | 12718 |
| Number of genes in orthogroups | 9,316 | 15,974 | 8,929 | 8,998 | 10,481 | 8,688 | 9,228 | 9,695 | 9,591 | 11,999 | 10,325 | 11070 |
| Number of unassigned genes | 935 | 3,272 | 952 | 1,037 | 1,534 | 1,014 | 4,348 | 1,175 | 2,402 | 1,579 | 2,489 | 1648 |
| Percentage of genes in orthogroups | 90.9 | 83.0 | 90.4 | 89.7 | 87.2 | 89.5 | 68.0 | 89.2 | 80.0 | 88.4 | 80.6 | 87.0 |
| Percentage of unassigned genes | 9.1 | 17.0 | 9.6 | 10.3 | 12.8 | 10.5 | 32.0 | 10.8 | 20.0 | 11.6 | 19.4 | 13.0 |
| Number of orthogroups containing species | 8,365 | 10,058 | 8,390 | 8,142 | 8,481 | 8,011 | 6,747 | 8,625 | 7,934 | 9,442 | 7,942 | 8416 |
| Percentage of orthogroups containing species | 51.8 | 62.2 | 51.9 | 50.4 | 52.5 | 49.6 | 41.8 | 53.4 | 49.1 | 58.4 | 49.1 | 52.1 |
| Number of species-specific orthogroups | 45 | 755 | 29 | 36 | 162 | 21 | 444 | 100 | 342 | 240 | 320 | 179 |
| Number of genes in species-specific orthogroups | 202 | 2,977 | 130 | 122 | 552 | 63 | 1,546 | 358 | 1,064 | 853 | 1,288 | 1168 |
| Percentage of genes in species-specific orthogroups | 2.0 | 15.5 | 1.3 | 1.2 | 4.6 | 0.6 | 11.4 | 3.3 | 8.9 | 6.3 | 10.1 | 9.2 |

Sign: *Acep= A. cephalotes,	Adis= A. disparis*,	*Amel= A. mellifera,	Aros=A. rosae, Cflo= C. floridanum,	Csol= C. solmsi	, Dmel= D. melanogaster,	Fari=* *Fopius arisanus,	Mcin= M. cingulum,	Nvit	= N. vitripennis, Tcas= T. castaneum,	Tper=* *T. pretiosum*
