## Supplementary Table 15 for "The Genome of the Wasp *Anastatus disparis* Reveals Energy Metabolism Adaptations for Extreme Aggression"

**Supplementary Table 15. Primer sequences for dsRNA synthesis**

| Gene name | Primer |
| --- | --- |
| *apoLp* | Forward : 5’- TAATACGACTCACTATAGGGAGA ATGGCACTACCACCCCGGCT-3’  Reverse: 5’- TAATACGACTCACTATAGGGAGAGTCTTGGCACCGGCATCACCA-3’ |
| *GFP* | Forward : 5’- TAATACGACTCACTATAGGGGGTGATGCTACATACGGAAAG -3’  Reverse: 5’- TAATACGACTCACTATAGGGTTGTTTGTCTGCCGTGAT-3’ |
