## Supplementary Table 16 for "The Genome of the Wasp *Anastatus disparis* Reveals Energy Metabolism Adaptations for Extreme Aggression"

| Gene name | Forward | Reverse |
| --- | --- | --- |
| *Tret1-like* | 5’-TTGCTGTGGCTGCTTCATTT-3’ | 5’-ACTGCGTAAACGTGAAGGTG-3’ |
| *AKR1A1b-like* | 5’-GGCGATGAAGATACAACGGT-3’ | 5’-TCTGGTCTGTTGCCGTATGA-3’ |
| *iPLA(2)* | 5’-CGAGACCGCTTACAGAGAGT-3’ | 5’-GCTGACCAAGCAGCTTCAAT-3’ |
| *agl-like* | 5’-TAGACCACCTGGACCACCTA-3’ | 5’-TGCCATCACCATCGCTATCT-3’ |
| *Vg3-like* | 5’-ACAAGAGGACGAGGTTGTGT-3’ | 5’-TTCACCTTGGTGTAGACGCT-3’ |
| *PLA2* | 5’-CGGACCTCATAGGTTAGCGT-3’ | 5’-CAATGGCTAAGGGTGAACGG-3’ |
| *FAS-like* | 5’-CGCTTATGCAGATGGCTGTT-3’ | 5’-CCAACTGCTGCCATAGAACC-3’ |
| *4CL-like* | 5’-ACCACATCTCGCCTTCTCAA-3’ | 5’-CCGGGAACCTTGACTACGAA-3’ |
| *SCAD* | 5’-TGTGCAAGTTGTGGCGTTAT-3’ | 5’-GCTTTAGCCATTGTTGATGCAG-3’ |
| *stp-like* | 5’-GAGCTTGCACAGATGCAGAA-3’ | 5’-TGGAAGACCATTCCAGCACA-3’ |
| *Lip 1* | 5’-TACTTATCGGGCCTGGCAAA-3’ | 5’-TACATCGCGTGTTGCCATTT-3’ |
| *ApoLp* | 5’-GGCAAACGTCTTGGTGAACT-3’ | 5’-TGATTTGAGGTCGGCTCCTT-3’ |
| *GP* | 5’-CGCTTGGTCTTGCTGCTTAT-3’ | 5’-ATTCAGGTCTGGCCTTCTCC-3’ |
| *ADΔ11* | 5’-TGGGTAAAGATTCATCGGACAC-3’ | 5’-AAGAAGCGCCAACCAATGTG-3’ |
| *GCDH-like* | 5’-CCCAGCAAGCGATCCTAATG-3’ | 5’-GCACCTTTCTCGCCAATCAT-3’ |
| *LPIN1* | 5’-ATTAATGCCGCAACCCTCAC-3’ | 5’-CCAAAGCGTACATGGAAGGG-3’ |
| *LRP4* | 5’-ACTGCACTGCGGAACAATTT-3’ | 5’-TGCACATCCGCTAAGCTTTC-3’ |
| *AKR1A1a-like* | 5’-CAGCAGAGAAAGCAGTGACA-3’ | 5’-ACATCTGAAGGTCGCATAGC-3’ |
| *ACBP4* | 5’-ATGTATAATATTTTTGCTT-3’ | 5’-TTAAGTTTCTTTAAATTCC-3’ |
| *POA3-like* | 5’-GGCCTCGTACTTCTCCCTTT-3’ | 5’-GAAGGAAGGCAGCATAAGCC-3’ |
| *EF1A* | 5’-ACCACGAAGCTCTCCAAGAA-3’ | 5’-AATCTGCAGCACCCTTAGGT-3’ |

**Supplementary Table 16. Primer pairs used for expression analysis using qRT-PCR**
